# Neurocognitive and psychiatric markers for addiction: Common vs. specific (endo)phenotypes for opiate and stimulant dependence

**DOI:** 10.1101/480970

**Authors:** Elizabeth C. Long, Radka Kaneva, Georgi Vasilev, F. Gerard Moeller, Jasmin Vassileva

## Abstract

**BACKGROUND:** The differential utility of neurocognitive impulsivity and externalizing/internalizing traits as putative addiction endophenotypes among individuals dependent on opiates vs. stimulants is unclear. The present study aims to determine: (1) whether neurocognitive impulsivity dimensions and externalizing/internalizing traits are correlated between siblings discordant for opiate and stimulant dependence; and (2) which of these associations are common across substances and which are substance-specific.

**METHOD:** Pearson correlations between individuals with ‘pure’ heroin and ‘pure’ amphetamine dependence and their unaffected biological siblings (*n* = 37 heroin sibling pairs; *n* = 30 amphetamine sibling pairs) were run on 10 neurocognitive measures, 6 externalizing measures, and 5 internalizing measures. Sibling pair effects were further examined using regression.

**RESULTS:** Siblings discordant for heroin dependence were significantly correlated on delay aversion on the Cambridge Gambling Task, risk-taking on the Balloon Analogue Risk Task, sensation seeking, and hopelessness. Siblings discordant for amphetamine dependence were significantly correlated on quality of decision-making on the Cambridge Gambling Task, discriminability on the Immediate Memory Task, commission errors on the Go/No-Go Task, trait impulsivity, ADHD, and anxiety sensitivity.

**CONCLUSIONS:** Dimensions of impulsivity and externalizing/internalizing traits appear to aggregate among siblings discordant for substance dependence. Risk-taking propensity, sensation seeking, and hopelessness were specific for heroin sibling pairs. Motor/action impulsivity and trait impulsivity were specific to amphetamine sibling pairs. Decisional/choice impulsivity was common across both heroin and amphetamine sibling pairs. These findings provide preliminary evidence for the utility of neurocognitive impulsivity and externalizing/internalizing traits as candidate endophenotypes for substance dependence in general and for substance-specific dependencies.

## 1. Introduction

Twin and family studies have consistently shown that substance use disorders (SUDs) are influenced by genetic factors, with heritabilities on the order of ~50 % (Merikangas and McClair, 2012). However, heritability estimates can differ dramatically depending on specific drug classes, with heroin showing some of the highest unique heritability (~70 %) relative to the much lower specific heritability of stimulants (~30 %) (Goldman et al., 2005; Tsuang et al., 1998). Further, the high prevalence of polysubstance dependence in studies conducted in North America is recognized as a key source of sample heterogeneity in SUDs (Agrawal et al., 2007), leading to conflicting findings in the literature reporting SUD heritabilities. This heterogeneity and the complexity of SUDs explains in part why molecular genetic efforts, such as genome-wide association studies (GWAS), have failed to locate specific genes and to account for the same amount of genetic variance as twin studies.

A recent conceptual approach that can help reduce the heterogeneity of SUD phenotypes and provide a framework for identifying general and specific influences on SUDs is the “endophenotype” approach (Fineberg et al., 2010; Frederick and Iacono, 2006; Gilmore et al., 2010; Gottesman and Gould, 2003). Endophenotypes are measurable traits, intermediate between the clinical phenotype and the disease-susceptibility genotype (Fineberg et al., 2010; Gottesman and Gould, 2003), thought to be genetically “simpler” than SUDs themselves. Neurocognitive functions are particularly suitable as endophenotypes and are more objective than self-report measures. As examples, there is evidence for executive function deficits as endophenotypes for ADHD (Gau and Shang, 2010; Rommelse et al., 2008); for processing speed, working memory, and facial memory as endophenotypes for bipolar disorder (Glahn et al., 2010); and for memory and emotion processing accuracy and speed of attention as endophenotypes for schizophrenia (Savitz et al., 2005).

Of the various neurocognitive functions implicated in SUDs, neurocognitive dimensions of impulsivity have received some of the strongest support as a candidate endophenotype for SUDs (Bickel, 2015; Frederick and Iacono, 2006; Kreek et al., 2005; MacKillop, 2013). Neurocognitive impulsivity is characterized by multiple dimensions that are typically measured with tasks falling into one of two categories (Winstanley et al., 2010): (1) *Decisional/Choice* impulsivity, which refers to the tendency to choose immediate but smaller rewards over delayed but larger rewards and may involve deficits in delaying gratification and exerting self-control (Fineberg et al., 2010), assessed with decision-making tasks involving various risk, reward, and delay contingencies (Fineberg et al., 2010; Hamilton et al., 2015b); and (2) *Motor/Action* impulsivity, which refers to the ability to refrain from inhibiting inappropriate behaviors, assessed with response inhibition tasks (Fineberg et al., 2014; Hamilton et al., 2015a). These impulsivity dimensions may differ in important ways between individuals who are dependent on different classes of drugs such as opiates and stimulants (Badiani et al., 2011; Ersche and Sahakian, 2007; Fernández-Serrano et al., 2011; George and Koob, 2010; Rogers et al., 1999; Verdejo-Garcia et al., 2007).

Of potential endophenotypic significance for SUDs are also traits and disorders within the externalizing and internalizing spectra, shown to be some of the most reliable risk factors for SUDs (Hussong et al., 2011; King et al., 2004; Krueger et al., 2002; Krueger et al., 2007). These traits are characterized by distinct neurocognitive profiles that most generally fall on a continuum between impulsivity and cognitive (over)control. Externalizing traits are characterized by neurocognitive deficits in impulse control, including: (1) increased reward sensitivity (Bava and Tapert, 2010; Huijbregts et al., 2008; Stout et al., 2004; Stout et al., 2005); (2) decreased loss aversion (Ahn et al., 2014; Passarotti et al., 2010; Stout et al., 2004; Stout et al., 2005); (3) increased delay discounting (Kirby, K. N. et al., 1999; Reynolds, 2006; Reynolds and Fields, 2012); and (4) decreased response inhibition (Castellanos-Ryan et al., 2016; Castellanos-Ryan et al., 2011; Castellanos-Ryan et al., 2014; Nigg, 2000). In contrast, internalizing traits have been associated with a neurocognitive profile characterized by: (1) reward & punishment processing abnormalities (Roiser and Sahakian, 2013; Smoski et al., 2008); (2) increased loss and risk aversion (Smoski et al., 2008); (3) decreased delay discounting (Lempert and Pizzagalli, 2010; Liu et al., 2012); (4) increased attentional lapses (Erickson et al., 2005); and (5) increased negative affect (Etkin et al., 2011; Mitterschiffthaler et al., 2008; Song et al., 2017; Surguladze et al., 2005; Williams et al., 1996).

Currently, it is unclear whether different dimensions of impulsivity and externalizing and internalizing traits/disorders have differential utility as putative endophenotypes of dependence among individuals dependent on opiates vs. those who are dependent on stimulants. We have the unique opportunity to examine this question and investigate the relationships between a large number of neurocognitive, externalizing, and internalizing phenotypes in a sibling-pair design via a research program we have developed in Bulgaria, where we have access to rare populations of ‘pure’ (i.e., mono-substance dependent) heroin and amphetamine users (Ahn et al., 2014; Ahn and Vassileva, 2016; Segalà et al., 2015; Vassileva et al., 2011; Vassileva et al., 2014; Vassileva et al., 2007; Wilson and Vassileva, 2018; Wilson and Vassileva, 2016).

We specifically aim to (1) Investigate whether discrete neurocognitive dimensions of impulsivity as well as symptoms of externalizing and internalizing traits/disorders are correlated between siblings discordant for opiate and stimulant dependence, thereby providing preliminary evidence for their utility as candidate endophenotypes for SUDs; and (2) Determine which of these putative endophenotypes are common and which are substance-specific. We expect neurocognitive dimensions of impulsivity, externalizing traits/disorders, and internalizing traits/disorders to be correlated among individuals with SUDs and their unaffected siblings, which can provide evidence for their utility as endophenotypes. Further, based on previous findings (Ahn and Vassileva, 2016), we expect some to be common across substances and others to be substance-specific.

## 2. Methods

### 2.1 Participants

Participants were selected from a larger study of impulsivity among individuals with heroin and amphetamine dependence in Sofia, Bulgaria. Participants were recruited via flyers placed in substance abuse clinics, nightclubs, bars, and cafes in Sofia, as well as by word of mouth. Initial screening for medical and substance use histories was done by telephone and on-site. Inclusion criteria were as follows: (1) age between 18 and 50 years; (2) estimated IQ > 75; (3) minimum of 8th grade education; (4) no history of neurological illness; (5) HIV seronegative status; and (6) negative breathalyzer test for alcohol and negative urine toxicology screen for amphetamines, methamphetamines, cocaine, opiates, methadone, cannabis, benzodiazepines, barbiturates, and MDMA.

In order to assess the utility of various personality, psychiatric, and neurocognitive measures as candidate endophenotypes for opiate and stimulant dependence, non-affected siblings of the participants with substance dependence were also recruited. Participants in the current study included 37 individuals with heroin dependence and 37 of their biological siblings with no history of substance dependence, as well as 30 individuals with amphetamine dependence and 30 of their non-dependent biological siblings. The majority of substance dependent individuals were in protracted abstinence (i.e. >1 year) at the time of testing. Critically, the majority of substance dependent individuals were mono-dependent on either heroin or amphetamines. There were a few participants with polysubstance dependence (9 in the heroin group and 7 in the amphetamine group), but these individuals were primarily dependent on the respective substance. The affected sibling with substance dependence was coded as ‘Sibling 1’, while the unaffected sibling was coded as ‘Sibling 2’.

### 2.2 Measures

Participants were administered 7 commonly used neurocognitive tasks measuring different dimensions of impulsivity from which 10 performance indices were selected, 6 measures of externalizing traits and disorders, and 5 measures of internalizing traits and disorders. Substance dependence was measured with the Substance Abuse module of the Structured Clinical Interview for DSM-IV (SCID-I; (First and Gibbon, 2004)).

#### 2.2.1 Neurocognitive tasks

While each of the 7 neurocognitive tasks generated several performance indices, we typically chose the most commonly used ones, as described below. For all measures except the Iowa Gambling Task and the Go/Stop Task, higher scores indicate higher levels of impulsivity.

##### 2.2.1.1 Decisional/Choice Impulsivity

###### Iowa Gambling Task

(IGT; (Bechara et al., 2000)). The IGT measures affective decision-making under conditions of ambiguity and uncertainty, which involves learning of rewards and punishment to guide decision-making. The total net score was used as the performance measure.

###### Cambridge Gambling Task

(CGT; (Rogers et al., 1999)). The CGT, part of the Cambridge Neuropsychological Test Automated Battery (CANTAB; (Robbins et al., 1994)), is a probabilistic task indexing decision-making and risk-taking outside of a learning context, where no uncertainty is involved. We used delay aversion (DA) and the quality of decision-making (QDM) as the performance measures.

###### Monetary Choice Questionnaire

(MCQ; (Kirby, Kris N et al., 1999)). The MCQ indexes delayed reward discounting (i.e., preference for smaller immediate rewards than larger delayed rewards). The *k* discounting rate parameter (log-transformed) and the mean number of inconsistent responses (INC) were used as indices of performance.

Unlike most other studies, we included INC because it provides additional information about choice consistency, which may be important to examine. Our group is currently testing novel computational models of delay discounting such as random utility models (Dai et al., 2016) that include choice consistency as a key parameter.

###### Balloon Analogue Risk Task

(BART; (Lejuez et al., 2002)). The BART measures risk preferences and tolerance for exposure to risk in pursuit to a reward. We used the pumps adjusted average as a performance measure.

##### 2.2.1.2 Motor/Action Impulsivity

###### Immediate Memory Task

(IMT; (Dougherty et al., 2003)). The IMT is a modified continuous performance task with complex demands on inhibitory control, working memory, and sustained attention. We selected two commonly used parameteric performance measures on the IMT: discriminability (*d’*) and response bias (β).

###### Stop Signal Task

(STOP; (Dougherty et al., 2003)). The task assesses the ability to withhold a prepotent response that has already been initiated. The performance measure used was the 150 msec inhibition, calculated by dividing the failures to inhibit a response by the correct detections after a stop signal appearing 150 ms after the appearance of the target.

###### Go/No-Go Task

(GNG; (Lane et al., 2007)). The GNG assesses the ability to inhibit prepotent responding. We used the number of commission errors as the performance measure.

#### 2.2.2 Externalizing traits/disorders

Unless otherwise noted, the total scores on each of the 6 measures were used for the analyses.

##### Barratt Impulsiveness Scale - 11^th^ revision

(BIS-11; (Patton et al., 1995)). The BIS is a 30-item self-report scale assessing common impulsive behaviors. Participants were instructed to indicate the extent to which they agree with each item, ranging from 1 (rarely/never) to 4 (almost always/always).

##### Sensation Seeking Scale - Version V

(SSS-V; (Zuckerman, 1996)). The SSS is a 40-item self-report scale reflecting a propensity to engage in novel, risky, or arousing types of behaviors. We used the existing (unpublished) Bulgarian version of the scale.

##### Buss-Warren Aggression Questionnaire

(BUSS; (Buss and Warren, 2000)). The BUSS is a 34-item self-report screening for aggression, a prominent behavioral manifestation of impulsivity. We used the Bulgarian version of the scale (Popov et al., 2016a).

##### Wender Utah Rating Scale

(WURS; (Ward et al., 1993)). The WURS is a self-report scale used to evaluate adults for childhood symptoms of ADHD. We used the recently validated 25-item Bulgarian version of the scale (Nedelchev et al., 2016).

##### Antisocial Personality Disorder (ASPD)

Symptom counts of antisocial personality disorder were obtained via the Antisocial Personality Module of the Structured Clinical Interview for DSM-IV (SCID-I; (First and Gibbon, 2004)).

##### Psychopathy Checklist, Screening Version

(PCL:SV; (Hart et al., 1995)). The PCL:SV is a 12-item, interviewer-completed scale based on a semi-structured interview that assesses interpersonal/affective and antisocial psychopathy (Hare, 1991). The Bulgarian adaptation of the instrument (Wilson et al., 2014) was used.

#### 2.2.3 Internalizing traits/disorders

Unless otherwise noted, the total scores on each of the 6 measures were used for the analyses.

##### Beck Depression Inventory - II

(BDI-II; (Beck et al., 1996)). The BDI-II is a 21- item scale that measures severity of depression symptoms during the last two weeks using a 4-point Likert Scale. We used the existing (unpublished) Bulgarian version of the scale.

##### Substance Use Risk Profile Scale, Hopelessness Subscale

(SURPS; (Woicik et al., 2009)). The SURPS is a 23-item self-report scale assessing 4 personality traits associated with increased risk for substance misuse on a 4-point Likert Scale (impulsivity, sensation seeking, hopelessness, and anxiety sensitivity). We only used the hopelessness subscale because the other subscales overlap with other measures already included. We used the recently validated Bulgarian version of the SURPS (Long et al., 2018).

##### Anxiety Sensitivity Index

(ANXSI; (Reiss et al., 1986)). The ANXSI is a 16-item, 5-point Likert scale that measures anxiety sensitivity as a global construct composed of several factors differentiating fear of specific anxiety symptoms and associated catastrophic consequences (Olthuis et al., 2014).

##### State-Trait Anxiety Inventory, State Anxiety subscale

(STAI-S; (Spielberger and Jacobs, 1983)). The STAI is a 20 item, 4-point Likert scale that consists of Trait and State subscales. Only the State subscale was used. We used the existing Bulgarian adaptation (Shtetinski and Paspalanov, 2008).

##### Toronto Alexithymia Scale – 20

(TAS-20; (Bagby et al., 1994)). The TAS is a 20- item scale designed to measure alexithymia associated with difficulties identifying and describing one’s own feelings (Leising et al., 2009). We used the recently translated and validated Bulgarian version (Popov et al., 2016b).

### Analyses

All analyses were conducted in R (R Development Core Team, 2013). Pearson correlations were run between individuals with heroin dependence and their non-dependent biological siblings, and between individuals with amphetamine dependence and their non-dependent biological siblings, on the 10 neurocognitive measures, 6 externalizing measures, and 5 internalizing measures, using the *corr.test* function.

To further examine the extent to which neurocognitive, externalizing, and internalizing dimensions in the affected sibling predict these traits in the unaffected sibling, we ran linear regressions using the *lm* function, with the independent variable being the dimensions for the affected sibling (sibling 1) and the dependent variable being the dimensions for the unaffected sibling (sibling 2). Separate models were run for each variable. Due to zero-inflation for ASPD, PCL:SV, and BDI-II, negative binomial regressions were run for these variables using the *glm.nb* function. Age and sex were included as covariates in all regression models. Directionality of the IGT and STOP was reversed for the correlations and regressions to maintain consistency across measures, such that high scores reflect high impulsivity on all measures.

## 3. Results

### 3.1 Descriptive statistics

The means and standard deviations (SDs) for the neurocognitive, externalizing, and internalizing measures are shown in Table 1, stratified by affected and unaffected siblings. For the neurocognitive measures, the means were fairly similar between the affected vs. unaffected siblings for both the discordant heroin and amphetamine pairs, with the exception of the BART, where the affected sibling demonstrated higher impulsivity. Opposite patterns of performance were observed between the discordant heroin pairs and discordant amphetamine pairs on the STOP Task and the GNG Task. For the discordant heroin pairs, the affected siblings scored higher on the STOP Task and the unaffected siblings scored higher on the GNG Task, whereas for the discordant amphetamine pairs the unaffected siblings scored higher on STOP Task and the affected siblings scored higher on the GNG Task. The unaffected siblings among the discordant amphetamine pairs also had a higher mean score on the IGT. Note that higher scores on the STOP Task and IGT indicate lower levels of impulsivity, whereas higher scores on all other measures indicate higher levels of impulsivity. For most of the externalizing and internalizing traits/disorders, the affected siblings had higher scores than the unaffected siblings, except for hopelessness among the discordant amphetamine pairs, where the siblings’ scores were similar.

**Table 1.** Descriptive statistics

|  | **Sibling Pairs Discordant for Heroin Dependence, N = 37** |  |
| --- | --- | --- |
|  | **Affected Sibling** | **Unaffected Sibling** |
| **Neurocognitive Measures, mean (SD)** |  |  |
| Iowa Gambling Task (IGT), Net Score | -0.66 (23.69) | -0.81 (29.78) |
| Cambridge Gambling Task (CGT), Delay aversion | 0.29 (0.21) | 0.37 (0.24) |
| Cambridge Gambling Task (CGT), Quality of decision making | 0.86 (0.15) | 0.88 (0.09) |
| Monetary Choice Questionnaire (MCQ), Log overall *k* | -1.48 (0.65) | -1.55 (0.68) |
| Monetary Choice Questionnaire (MCQ), # of inconsistencies | 0.21 (0.42) | 0.62 (1.09) |
| Balloon Analogue Risk Task (BART), Pumps adjusted average | 42.20 (12.48) | 37.52 (14.76) |
| Immediate Memory Task (IMT), Discriminability (*d'*) | 1.32 (0.54) | 1.29 (0.43) |
| Immediate Memory Task (IMT), Response bias ($\beta$) | 0.75 (0.43) | 0.84 (0.43) |
| Go/Stop Task (STOP), 150 msec inhibition | 73.57 (17.13) | 70.68 (20.89) |
| Go/No-Go Task (GNG), Commission errors | 13.44 (8.45) | 16.59 (11.94) |
| **Externalizing Traits/Disorders, mean (SD)** |  |  |
| Barratt Impulsiveness Scale-11 (BIS-11) | 61.67 (10.40) | 56.03 (7.61) |
| Sensation Seeking Scale (SSS-V) | 18.83 (7.16) | 17.49 (7.34) |
| Buss Warren Aggression Questionnaire (BUSS) | 44.42 (9.28) | 38.03 (9.33) |
| Wender Utah Rating Scale for ADHD (WURS) | 37.54 (19.64) | 20.16 (11.47) |
| Antisocial Personality Disorder (ASPD), # of symptoms | 3.78 (1.89) | 0.19 (0.57) |
| Psychopathy Checklist (PCL:SV) | 12.56 (4.99) | 2.54 (2.57) |
| **Internalizing Traits/Disorders, mean (SD)** |  |  |
| Beck Depression Inventory II (BDI-II) | 7.31 (5.31) | 6.30 (7.13) |
| Hopelessness, Substance Use Risk Profile (SURPS) | 12.87 (3.96) | 12.61 (3.59) |
| Anxiety Sensitivity Index (ANXSI) | 19.35 (9.46) | 17.05 (8.24) |
| State Anxiety (STAI) | 33.78 (7.23) | 32.35 (10.15) |
| Toronto Alexithymia Scale (TAS) | 47.07 (10.72) | 44.32 (9.66) |
|  | **Sibling Pairs Discordant for Amphetamine Dependence, N = 30** |  |
|  | **Affected Sibling** | **Unaffected Sibling** |
| **Neurocognitive Measures, mean (SD)** |  |  |
| Iowa Gambling Task (IGT), Total Net Score | 0.72 (26.53) | 5.66 (27.59) |
| Cambridge Gambling Task (CGT), Delay aversion | 0.31 (0.18) | 0.25 (0.19) |
| Cambridge Gambling Task (CGT), Quality of decision making | 0.89 (0.08) | 0.88 (0.19) |
| Monetary Choice Questionnaire (MCQ), Log overall *k* | -1.49 (0.53) | -1.68 (0.61) |
| Monetary Choice Questionnaire (MCQ), # of inconsistencies | 0.33 (0.61) | 0.63 (1.19) |
| Balloon Analogue Risk Task (BART), Pumps adjusted average | 45.49 (13.12) | 42.50 (12.94) |
| Immediate Memory Task (IMT), Discriminability (*d'*) | 1.00 (0.50) | 1.14 (0.48) |
| Immediate Memory Task (IMT), Response bias ($\beta$) | 0.74 (0.23) | 0.85 (0.29) |
| Go/Stop Task (STOP), 150 msec inhibition | 64.00 (18.86) | 68.67 (18.84) |
| Go/No-Go Task (GNG), Commission errors | 17.37 (9.70) | 15.76 (8.12) |
| **Externalizing Traits/Disorders, mean (SD)** |  |  |
| Barratt Impulsiveness Scale-11 (BIS-11) | 67.20 (10.79) | 59.90 (11.04) |
| Sensation Seeking Scale V (SSS-V) | 23.70 (5.99) | 17.23 (6.89) |
| Buss Warren Aggression Questionnaire (BUSS) | 45.50 (13.40) | 37.57 (10.37) |
| Wender Utah Rating Scale for ADHD (WURS) | 37.93 (14.12) | 22.83 (14.17) |
| Antisocial Personality Disorder (ASPD), # of symptoms | 2.73 (1.66) | 0.67 (1.09) |
| Psychopathy Checklist (PCL:SV) | 10.87 (5.08) | 3.10 (3.46) |
| **Internalizing Traits/Disorders, mean (SD)** |  |  |
| Beck Depression Inventory II (BDI-II) | 7.17 (3.92) | 6.53 (5.99) |
| Hopelessness, Substance Use Risk Profile (SURPS) | 11.43 (2.83) | 11.60 (3.08) |
| Anxiety Sensitivity Index (ANXSI) | 21.30 (0.44) | 17.00 (9.60) |
| State Anxiety (STAI) | 32.40 (5.37) | 31.83 (7.92) |
| Toronto Alexithymia Scale (TAS) | 46.24 (8.93) | 42.60 (10.28) |

### 3.2 Correlations between siblings discordant for SUDs

The significant correlations between siblings discordant for SUDs on neurocognitive, externalizing, and internalizing measures are listed in Table 2 (please see Supplemental Tables 1, 2, and 3 for the full correlation matrices). The within-pair, within-trait (i.e., the correlations between sibling pairs on the same trait) correlations are shown in the top of the table, whereas the within-pair, cross-trait correlations (i.e., the correlations between sibling pairs on different traits) are shown in the bottom of the table.)

**Table 2.**
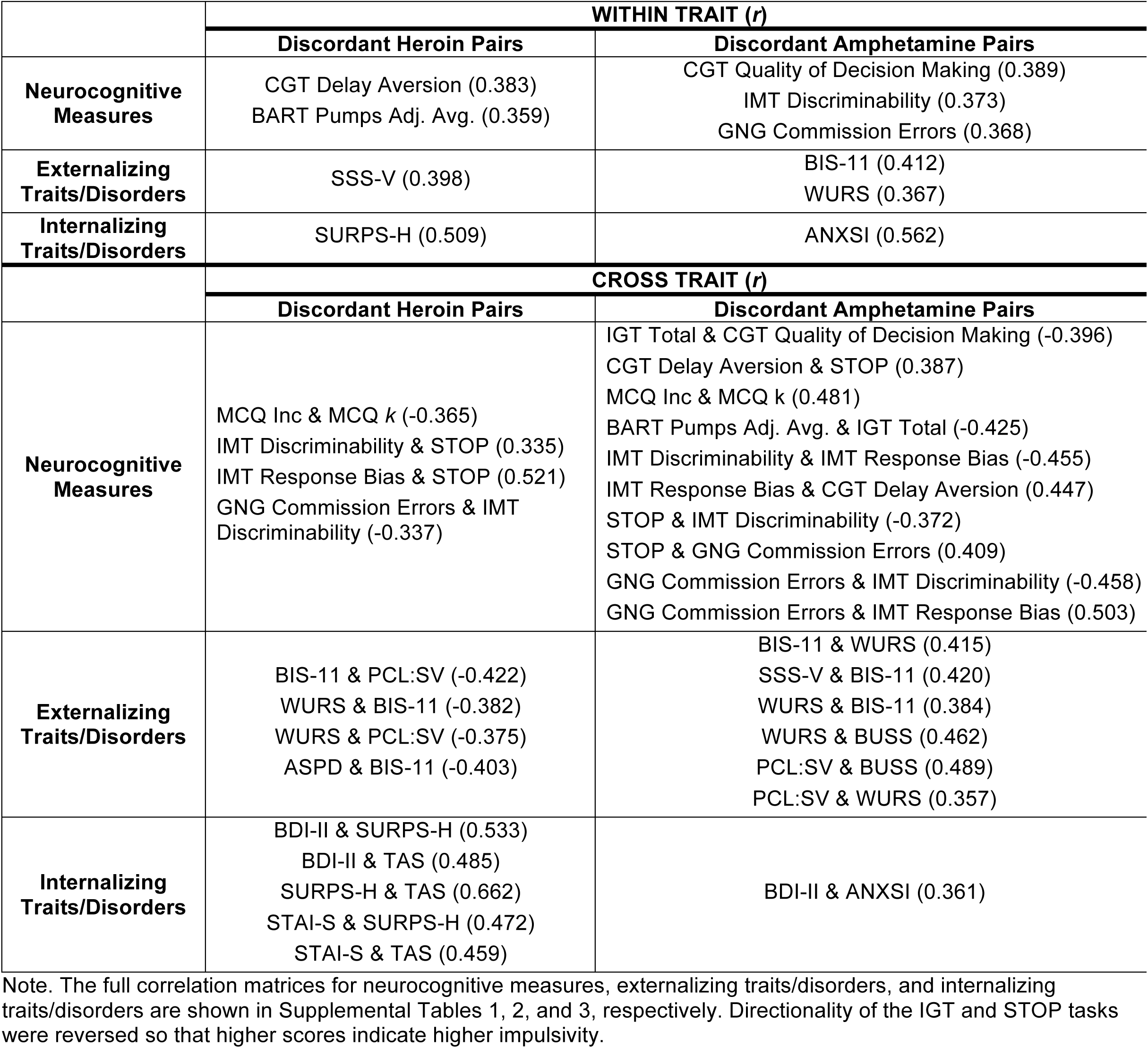
Significant within and cross trait correlations between siblings discordant for SUDs

**Table 3.** Sibling effects - Regression results

|  | **Sibling Pairs Discordant for Heroin Dependence, N = 37** |  |
| --- | --- | --- |
| **Neurocognitive Measures** | **Beta (SE)** | **p-value** |
| Iowa Gambling Task (IGT), Net Score* | -0.147 (0.219) | 0.509 |
| Cambridge Gambling Task (CGT), Delay aversion | 0.436 (0.219) | 0.059 |
| Cambridge Gambling Task (CGT), Quality of decision making | 0.061 (0.117) | 0.608 |
| Monetary Choice Questionnaire (MCQ), Log overall *k* | -0.029 (0.197) | 0.883 |
| Monetary Choice Questionnaire (MCQ), # of inconsistencies | -0.145 (0.540) | 0.790 |
| Balloon Analogue Risk Task (BART), Pumps adjusted average | **0.431 (0.201)** | **0.040** |
| Immediate Memory Task (IMT), Discriminability (*d'*) | 0.181 (0.144) | 0.217 |
| Immediate Memory Task (IMT), Response bias ($\beta$) | -0.086 (0.197) | 0.664 |
| Go/Stop Task (STOP), 150 msec inhibition* | 0.013 (0.214) | 0.952 |
| Go/No-Go Task (GNG), Commission errors | 0.003 (0.241) | 0.990 |
| **Externalizing Traits/Disorders** |  |  |
| Barratt Impulsiveness Scale-11 (BIS-11) | 0.089 (0.129) | 0.498 |
| Sensation Seeking Scale (SSS-V) | **0.397 (0.165)** | **0.022** |
| Buss Warren Aggression Questionnaire (BUSS) | 0.019 (0.179) | 0.916 |
| Wender Utah Rating Scale for ADHD (WURS) | -0.050 (0.107) | 0.641 |
| Antisocial Personality Disorder (ASPD), # of symptoms** | -0.401 (0.288) | 0.164 |
| Psychopathy Checklist (PCL:SV)** | 0.001 (0.038) | 0.971 |
| **Internalizing Traits/Disorders** |  |  |
| Beck Depression Inventory II (BDI-II)** | 0.031 (0.037) | 0.400 |
| Substance Use Risk Profile (SURPS), Hopelessness subscale | **0.448 (0.188)** | **0.028** |
| Anxiety Sensitivity Index (ANXSI) | 0.008 (0.152) | 0.956 |
| State-Trait Anxiety Inventory (STAI), State Anxiety subscale | 0.000 (0.283) | 1.000 |
| Toronto Alexithymia Scale (TAS) | 0.332 (0.165) | 0.056 |
|  | **Sibling Pairs Discordant for Amphetamine Dependence, N = 30** |  |
|  | **Beta** | **p-value** |
| **Neurocognitive Measures** |  |  |
| Iowa Gambling Task (IGT), Total* | -0.016 (0.207) | 0.938 |
| Cambridge Gambling Task (CGT), Delay aversion | -0.196 (0.220) | 0.383 |
| Cambridge Gambling Task (CGT), Quality of decision making | 0.838 (0.446) | 0.074 |
| Monetary Choice Questionnaire (MCQ), Log overall *k* | -0.036 (0.233) | 0.879 |
| Monetary Choice Questionnaire (MCQ), # of inconsistencies | 0.169 (0.389) | 0.669 |
| Balloon Analogue Risk Task (BART), Pumps adjusted average | 0.266 (0.187) | 0.168 |
| Immediate Memory Task (IMT), Discriminability (*d'*) | 0.327 (0.175) | 0.073 |
| Immediate Memory Task (IMT), Response bias ($\beta$) | 0.115 (0.243) | 0.639 |
| Go/Stop Task (STOP), 150 msec inhibition* | 0.131 (0.199) | 0.514 |
| Go/No-Go Task (GNG), Commission errors | 0.314 (0.158) | 0.059 |
| **Externalizing Traits/Disorders** |  |  |
| Barratt Impulsiveness Scale 11 (BIS) | **0.448 (0.191)** | **0.027** |
| Sensation Seeking Scale V (SSS) | -0.129 (0.209) | 0.541 |
| Buss Warren Aggression Questionnaire (BUSS) | 0.220 (0.146) | 0.143 |
| Wender Utah Rating Scale for ADHD (WURS) | **0.393 (0.183)** | **0.041** |
| Antisocial Personality Disorder (ASPD), # of symptoms** | -0.010 (0.217) | 0.963 |
| Psychopathy Checklist (PCL:SV)** | 0.058 (0.052) | 0.263 |
| **Internalizing Traits/Disorders** |  |  |
| Beck Depression Inventory II (BDI-II)** | 0.036 (0.050) | 0.468 |
| Substance Use Risk Profile (SURPS), Hopelessness subscale | -0.124 (0.254) | 0.632 |
| Anxiety Sensitivity Index (ANXSI) | **0.469 (0.136)** | **0.002** |
| State-Trait Anxiety Inventory (STAI), State Anxiety subscale | -0.434 (0.299) | 0.159 |
| Toronto Alexithymia Scale (TAS) | -0.198 (0.226) | 0.388 |

Note. \*The direction of effect was reversed for the IGT and STOP variables so that higher scores reflect higher impulsivity, to be consistent with all other measures. \*\*Negative binomial regressions were run for ASPD, PCL:SV, and BDI due to zero-inflation. All models included age and sex as covariates.

#### 3.2.1 Neurocognitive measures

Among sibling pairs discordant for *heroin dependence*, there were significant positive within-trait sibling pair correlations on two measures of decisional/choice impulsivity: delay aversion on the CGT and pumps adjusted average on the BART. There were significant cross-trait positive correlations between discriminability on the IMT and inhibition on the STOP Task and between response bias on the IMT and inhibition on the STOP Task. There were significant cross-trait negative correlations between the discounting rate parameter (*k*) on the MCQ and number of inconsistencies on the MCQ, and between commission errors on the GNG Task and discriminability on the IMT.

Among sibling pairs discordant for *amphetamine dependence*, there were significant within-trait positive correlations on two measures of motor/action impulsivity: discriminability on the IMT and commission errors on the GNG Task. There was also a significant within-trait positive correlation on a measure of decisional/choice impulsivity, quality of decision-making on the CGT. There were a number of significant cross-trait positive correlations: delay aversion on the CGT and inhibition on the STOP Task; discounting rate parameter (*k*) on the MCQ and number of inconsistencies on the MCQ; response bias on the IMT and delay aversion on the CGT; inhibition on the STOP Task and commission errors on the GNG Task; and commission errors on the GNG Task and response bias on the IMT. There were also a number of significant cross-trait negative correlations: overall net score on the IGT and quality of decision making on the CGT; pumps adjusted average on the BART and overall net score on the IGT; discriminability on the IMT and response bias on the IMT; inhibition on the STOP Task and discriminability on the IMT; and commission errors on the GNG Task and discriminability on the IMT.

#### 3.2.2 Externalizing traits/disorders

The sibling pairs discordant for *heroin dependence* were significantly and positively correlated on sensation seeking (SSS-V). There were no significant cross-trait positive correlations, but there were a number of significant cross-trait negative correlations between trait impulsivity (BIS-11) and psychopathy (PCL:SV); ADHD (WURS) and trait impulsivity (BIS-11); ADHD (WURS) and psychopathy (PCL:SV); and ASPD and trait impulsivity (BIS-11).

Among the sibling pairs discordant for *amphetamine dependence* there were significant within-trait positive correlations on trait impulsivity (BIS-11) and ADHD (WURS). There were a number of significant cross-trait positive correlations between sensation seeking (SSS-V) and trait impulsivity (BIS-11); ADHD (WURS) and trait impulsivity (BIS-11); ADHD (WURS) and aggression (BUSS); psychopathy (PCL:SV) and aggression (BUSS); and psychopathy (PCL:SV) and ADHD (WURS). There were no significant cross-trait negative correlations, which is the exact opposite pattern to the one we found with the siblings discordant for heroin dependence.

#### 3.2.3 Internalizing Traits/Disorders

The sibling pairs discordant for heroin dependence were significantly and positively correlated on the Hopelessness subscale of the SURPS. There were a number of significant and positive cross-trait correlations: depression (BDI-II) and hopelessness (SURPS); depression (BDI-II) and alexithymia (TAS-20); hopelessness (SURPS) and alexithymia (TAS-20); state anxiety (STAI-S) and hopelessness (SURPS); and state anxiety (STAI-S) and alexithymia (TAS-20). There were no significant cross-trait negative correlations.

The sibling pairs discordant for amphetamine dependence were significantly and positively correlated on the Anxiety Sensitivity Index (ANXSI). There was only one significant cross-trait positive correlation between depression (BDI-II) and anxiety sensitivity (ANXSI). There were no significant cross-trait negative correlations.

### 3.3 Regressions

Finally, we examined the extent to which neurocognitive, externalizing, and internalizing dimensions in the affected sibling predict these dimensions in the unaffected sibling with linear and negative binomial regressions. These results are presented in Table 3. The patterns that were observed for the within-trait correlations were generally reflected in the regression results. For example, risk-taking (indexed by the BART), sensation seeking, and hopelessness in the siblings with heroin dependence significantly predicted these traits in the unaffected siblings. Although significantly correlated, delay aversion on the CGT did not reach statistical significance in the regression, though it was trending (*p* = 0.059).

Likewise, trait impulsivity (BIS-11), ADHD (WURS), and anxiety sensitivity (ANXSI) in the siblings with amphetamine dependence significantly predicted these traits in the unaffected siblings. However, the neurocognitive measures that were significantly correlated did not reach statistical significance (quality of decision making on the CGT, discriminability on the IMT, and commission errors on the GNG Task), but were similarly trending (*p* = 0.074, *p* = 0.073, and *p* = 0.059, respectively).

## 4. Discussion

Using a sample of Bulgarian individuals with ‘pure’ heroin and amphetamine dependence and their unaffected siblings, we investigated sibling pair correlations on neurocognitive dimensions of impulsivity, externalizing traits/disorders, and internalizing traits/disorders in order to explore their potential utility as common vs. specific endophenotypes for opiate and stimulant dependence. Results revealed both common and substance-specific associations. Decisional/choice impulsivity was common across both heroin and amphetamine sibling pairs (delay aversion on the CGT and BART for discordant heroin pairs; quality of decision making on the CGT for discordant amphetamine pairs), whereas motor/action impulsivity (discriminability on the IMT and commission errors on the GNG) was specific to amphetamine sibling pairs. Sensation seeking (SSS-V) and hopelessness (SURPS-H) were specific to discordant heroin sibling pairs, whereas trait impulsivity (BIS-11), ADHD (WURS), and anxiety sensitivity (ANXSI) were specific to discordant amphetamine sibling pairs. These results are consistent with previous literature showing that impulsivity and anxious-impulsive personality traits, but not sensation seeking, are candidate endophenotypes for stimulant dependence (Ersche et al., 2012; Ersche et al., 2010).

Of interest is the opposite direction of effects for some of the cross-trait correlations among the discordant heroin sibling pairs compared to the discordant amphetamine sibling pairs. The correlation between the delay discounting parameter *k* and the number of inconsistencies on the MCQ was positive for the discordant heroin pairs, but negative for the discordant amphetamine pairs. This pattern of results suggests that the discordant heroin pairs performed similarly on these measures, whereas the discordant amphetamine pairs differed in their performance, highlighting the value of adding the inconsistency measure. We believe this measure may offer useful information, as evidenced by the current results, and are currently testing novel computational models of delay discounting which include choice variability as a key parameter (Kvam et al., 2018).

Additionally, all the significant cross-trait correlations for the externalizing traits were negative for the discordant heroin pairs, but positive for the discordant amphetamine pairs. There were also many more significant cross-trait correlations for the internalizing traits among the discordant heroin pairs relative to the discordant amphetamine pairs. Together, these results suggest that the externalizing spectrum aggregates in siblings discordant for amphetamine dependence, whereas the internalizing spectrum aggregates in siblings discordant for heroin dependence. This is in line with previous work from our group, which similarly shows that internalizing traits such as depression and anxiety are significant predictors of heroin but not amphetamine dependence, whereas externalizing traits such as disinhibited sensation seeking and hostility predict amphetamine but not heroin dependence (Ahn and Vassileva, 2016).

One potential explanation for why internalizing traits may be endophenotypes specific to heroin dependence may be related to the notion of “+ hyperkatifeia,” which refers to increases in emotional distress and emotional pain experienced during withdrawal and abstinence from chronic drug use (Shurman et al., 2010). It is possible that individuals who are genetically predisposed to internalizing disorders are more likely to find the analgesic effects of opiates more reinforcing than individuals who are genetically predisposed to externalizing disorders.

Finally, our findings highlight the utility of simultaneously examining multiple neurocognitive, externalizing, and internalizing dimensions. Research indicates that the most powerful genetic approaches often involve multivariate, rather than univariate analyses of individual characteristics and traits (Iacono et al., 2018). Similarly, it has been noted that examination of multivariate candidate (endo)phenotypes may increase the power to detect genetic effects (Van Der Sluis et al., 2010). One pervasive problem in genetic association studies is the “missing heritability” problem, namely that the variance explained by genetic variants from GWAS studies is very small compared to the heritability estimates obtained from family studies (Van Der Sluis et al., 2010). While genetic heterogeneity is often invoked as an explanation, the manner in which complex phenotypic traits are measured and modeled are equally important contributors to the “missing heritability” problem but have received much less attention in the literature. Despite the multidimensionality of traits measured by psychometric, diagnostic, and neurocognitive instruments, most GWAS studies typically use total sum scores that do not reflect the underlying phenotypic multidimensionality. The current findings may inform future multivariate multilevel models of complex phenotypes and increase understanding of the complex relationship between multiple neurocognitive and personality phenotypes that may help redefine putative endophenotypes as multi-level combination of measures (Bilder et al., 2009), the next critical step in the endophenotype approach (Sabb et al., 2009).

## Limitations

Despite clear strengths of the present study (e.g., use of rare individuals with “pure” dependencies; comprehensive assessment of impulsivity, externalizing and internalizing dimensions, inclusion of non-affected biological siblings), our findings should be considered within the context of two limitations. First, our sample size was small. Second, our sample consisted entirely of individuals from Bulgaria. Although this limitation was necessary to permit use of individuals with pure dependencies, it is unclear if our findings will generalize to other populations.

## Conclusion

Neurocognitive dimensions of impulsivity, externalizing, and internalizing traits appear to aggregate differentially among siblings discordant for heroin and amphetamine dependence. These findings provide preliminary evidence for the utility of these traits as common and specific candidate endophenotypes for opiate and stimulant dependence.

## Supporting information

