## Supplementary material for "Neurocognitive and psychiatric markers for addiction: Common vs. specific (endo)phenotypes for opiate and stimulant dependence"

**Supplemental Table 1.** Correlations between siblings discordant for SUDs on neurocognitive measures

| | Sib 1 IGT | Sib 1 CGT, DA | Sib 1 CGT, QDM | Sib 1 MCQ, *k* | Sib 1 MCQ, INC | Sib 1 BART | Sib 1 IMT, D | Sib 1 IMT, B | Sib 1 STOP | Sib 1 GNG |
|---|---|---|---|---|---|---|---|---|---|---|
| **Sibling Pairs Discordant for Heroin Dependence** | | | | | | | | | | |
| Sib 2 IGT, Total* | -0.101 | -0.070 | -0.082 | -0.045 | -0.170 | -0.033 | -0.194 | 0.090 | -0.246 | -0.002 |
| Sib 2 CGT, Delay aversion (DA) | 0.190 | **0.383** | -0.357 | -0.199 | -0.049 | -0.169 | 0.040 | 0.225 | -0.018 | 0.105 |
| Sib 2 CGT, Quality of decision making (QDM) | 0.076 | -0.112 | 0.063 | -0.049 | -0.163 | 0.139 | -0.194 | -0.056 | -0.198 | -0.092 |
| Sib 2 MCQ, Log overall *k* | 0.145 | 0.089 | -0.065 | -0.036 | 0.269 | -0.286 | -0.056 | -0.001 | 0.133 | 0.304 |
| Sib 2 MCQ, # of inconsistencies (INC) | 0.038 | 0.005 | -0.347 | **-0.365** | -0.058 | 0.104 | 0.148 | -0.250 | 0.011 | -0.064 |
| Sib 2 BART, Pumps adj. average | 0.168 | 0.349 | -0.214 | 0.204 | 0.010 | **0.359** | -0.053 | 0.195 | 0.081 | 0.059 |
| Sib 2 IMT, Discriminability (*d'*) | 0.088 | 0.011 | 0.122 | 0.029 | -0.159 | 0.059 | 0.216 | 0.139 | **0.335** | 0.047 |
| Sib 2 IMT, Response bias (β) | 0.073 | -0.009 | 0.084 | 0.113 | 0.013 | 0.241 | 0.179 | -0.002 | **0.521** | 0.143 |
| Sib 2 STOP, 150 msec inhibition* | 0.010 | 0.025 | 0.148 | 0.140 | -0.006 | -0.011 | 0.132 | 0.105 | 0.001 | -0.035 |
| Sib 2 GNG, Commission errors | -0.186 | 0.012 | 0.062 | -0.156 | 0.179 | -0.030 | **-0.337** | 0.080 | -0.261 | -0.012 |
| **Sibling Pairs Discordant for Amphetamine Dependence** | | | | | | | | | | |
| Sib 2 IGT, Total* | -0.003 | -0.097 | **-0.396** | 0.146 | -0.260 | -0.190 | 0.210 | -0.029 | 0.208 | 0.038 |
| Sib 2 CGT, Delay aversion (DA) | 0.157 | -0.129 | -0.059 | -0.302 | -0.047 | -0.077 | -0.057 | 0.284 | **0.387** | 0.028 |
| Sib 2 CGT, Quality of decision making (DM) | -0.036 | 0.098 | **0.389** | -0.143 | 0.242 | 0.098 | -0.011 | -0.006 | -0.124 | 0.360 |
| Sib 2 MCQ, Log overall *k* | 0.027 | 0.004 | 0.019 | 0.014 | -0.171 | -0.120 | -0.126 | 0.109 | 0.260 | 0.208 |
| Sib 2 MCQ, # of inconsistencies (INC) | -0.305 | 0.169 | -0.067 | **0.481** | 0.080 | -0.029 | -0.344 | 0.139 | 0.286 | 0.251 |
| Sib 2 BART, Pumps adj. average | **-0.425** | -0.067 | 0.029 | -0.260 | 0.017 | 0.230 | 0.100 | 0.073 | -0.069 | -0.270 |
| Sib 2 IMT, Discriminability (*d'*) | 0.194 | 0.096 | 0.288 | -0.078 | -0.260 | -0.033 | **0.373** | **-0.455** | -0.154 | -0.003 |
| Sib 2 IMT, Response bias (β) | -0.220 | **0.447** | -0.178 | 0.319 | 0.220 | 0.217 | -0.348 | 0.171 | 0.139 | 0.050 |
| Sib 2 STOP, 150 msec inhibition* | -0.003 | -0.206 | -0.073 | -0.206 | 0.035 | 0.134 | **-0.372** | 0.250 | 0.132 | **0.409** |
| Sib 2 GNG, Commission errors | 0.099 | -0.272 | 0.095 | -0.357 | -0.076 | 0.153 | **-0.458** | **0.503** | 0.150 | **0.368** |

**Note.** Bold = $p \leq 0.05$. The direction of effect was reversed for the IGT and STOP variables so that higher scores reflect higher impulsivity, to be consistent with all other measures.

**Supplemental Table 2.** Correlations between siblings discordant for SUDs on externalizing traits/disorders

| **Sibling Pairs Discordant for Heroin Dependence** | | | | | | |
|---|---|---|---|---|---|---|
| | Sib 1 BIS-11 | Sib 1 SSS-V | Sib 1 BUSS | Sib 1 WURS | Sib 1 ASPD | Sib 1 PCL |
| Sib 2 BIS-11, Total | 0.121 | 0.126 | -0.067 | -0.045 | -0.088 | **-0.422** |
| Sib 2 SSS-V, Total | 0.232 | **0.398** | -0.157 | 0.120 | -0.123 | -0.167 |
| Sib 2 BUSS, Total | -0.078 | 0.005 | 0.034 | 0.168 | -0.088 | -0.281 |
| Sib 2 WURS, Total | **-0.382** | -0.128 | -0.305 | -0.081 | -0.311 | **-0.375** |
| Sib 2 ASPD, # of symptoms | **-0.403** | -0.165 | -0.309 | 0.045 | -0.168 | -0.009 |
| Sib 2 PCL:SV, Total | -0.076 | 0.004 | -0.211 | 0.135 | 0.088 | 0.036 |
| **Sibling Pairs Discordant for Amphetamine Dependence** | | | | | | |
| Sib 2 BIS-11, Total | **0.412** | -0.048 | 0.335 | **0.415** | -0.248 | -0.147 |
| Sib 2 SSS-V, Total | **0.420** | -0.033 | 0.172 | 0.338 | -0.287 | -0.173 |
| Sib 2 BUSS, Total | 0.185 | -0.297 | 0.287 | 0.312 | -0.207 | -0.177 |
| Sib 2 WURS, Total | **0.384** | 0.011 | **0.462** | **0.367** | -0.005 | 0.032 |
| Sib 2 ASPD, # of symptoms | 0.216 | -0.100 | 0.210 | 0.197 | -0.032 | 0.041 |
| Sib 2 PCL:SV, Total | 0.325 | -0.132 | **0.489** | **0.357** | 0.029 | 0.109 |

**Note.** Bold = $p \leq 0.05$.

**Supplemental Table 3.** Correlations between siblings discordant for SUDs on internalizing traits/disorders

| | Sib 1 BDI-II | Sib 1 SURPS, H | Sib 1 ANXSI | Sib 1 STAI, S | Sib 1 TAS |
|---|---|---|---|---|---|
| **Sibling Pairs Discordant for Heroin Dependence** | | | | | |
| Sib 2 BDI-II, Total | 0.240 | **0.533** | 0.246 | 0.136 | **0.485** |
| Sib 2 SURPS, Hopelessness (H) | 0.316 | **0.509** | 0.069 | 0.034 | **0.662** |
| Sib 2 ANXSI, Total | -0.179 | -0.099 | 0.046 | -0.233 | 0.201 |
| Sib 2 STAI, State (S) | 0.231 | **0.472** | -0.174 | 0.042 | **0.459** |
| Sib 2 TAS, Total | -0.158 | 0.215 | 0.001 | -0.071 | 0.223 |
| **Sibling Pairs Discordant for Amphetamine Dependence** | | | | | |
| Sib 2 BDI-II, Total | 0.152 | -0.158 | **0.361** | -0.328 | 0.208 |
| Sib 2 SURPS, Hopelessness (H) | -0.214 | -0.027 | 0.139 | -0.175 | 0.018 |
| Sib 2 ANXSI, Total | 0.174 | -0.081 | **0.562** | -0.253 | 0.131 |
| Sib 2 STAI, State (S) | -0.041 | 0.119 | 0.129 | -0.305 | 0.161 |
| Sib 2 TAS, Total | -0.101 | -0.131 | 0.232 | -0.240 | -0.193 |

Note. Bold = $p \leq 0.05$.
